# Sex-specific fish recombination landscapes link recombination and karyotype evolution

**DOI:** 10.1101/2024.12.23.630081

**Authors:** Teemu Kivioja, Pasi Rastas

## Abstract

Meiotic recombination is an ubiquitous feature of sexual reproduction across eukaryotes. While recombination has been widely studied both theoretically and experimentally, the causes of its variation across species are still poorly understood. Composing a coherent view across species has been difficult because of the differences in recombination map generation and reporting of the results. Thus, fundamental questions like why recombination rates differ between sexes (heterochiasmy) in many but not all species remain unanswered. Here we present the first collection of recombination maps that allows quantitative comparisons across a diverse set of species. We generated sex-specific high-density linkage maps for 40 fish species using the same computational pipeline. Comparing the maps revealed that the higher genome-wide recombination rate in females compared to males was linked to the karyotype of the species. The difference between the sexes in the positioning of the crossovers was also highly variable and unrelated to the difference in their total number. Especially in males, CpG content of the sequence was a strong indicator of the broad scale distribution of crossovers between and within chromosomes. More generally, the collection of recombination landscapes can serve as a link between the theoretical and experimental work on recombination.

## Introduction

Meiotic recombination is a highly conserved process among eukaryotes. During meiosis, most sexually reproducing organisms require at least one recombination per homologous chromosome pair i.e. a crossover event that leads to reciprocal exchange of genetic material between them (we will use the term recombination to mean this specific process) [34]. On the other hand, each chromosome pair typically has only few crossovers irrespective of its size. Thus, as different eukaryotic groups have vastly different genome sizes, the average recombination rates vary by several orders of magnitude between distant groups [52]. Within these broad constraints, the positions and numbers of crossovers vary between species, populations, and even individuals [21, 34]. Which forces on one hand maintain the conserved, and on the other hand drive the variable aspects of recombination, remains poorly understood.

Recombination rate varies also between and within chromosomes of one species. First, as the number of crossovers per chromosome typically varies less than the chromosome size, the shortest chromosomes must have higher recombination rates than the longest ones. Second, in many vertebrates, including many but not all mammals, crossovers are directed to narrow (kilobase-scale) regions called hotspots determined by sequence-specific binding of the protein PRDM9 [16]. However, the complete PRMD9 needed for directing recombinations has been lost independently in many vertebrate lineages [4]. Instead, stable CpG-rich hotspots have been reported for many of these species such as dog [3] and birds [42] and recently some species have even been reported to have both types of hotspots [19, 22]. Third, even when hotspots are locally directed by the sequence-specificity of PRDM9 as in mouse, recombination appears to be organized also on other levels such as high-level chromatin structure [33]. Thus understanding recombination needs accurate and precise recombination landscapes that show the variation of recombination rate along each chromosome [34].

Interestingly, recombination rates also often differ between the sexes of the same species, a situation called heterochiasmy [39]. The sexes may have different genomewide recombination rates (we use the term heterochiasmy exclusively to mean that) but also the positioning of the crossovers along chromosome can be drastically different [39]. Numerous evolutionary hypotheses has been suggested to explain heterochiasmy but none of them has yet gained wide empirical support [8, 21, 25, 39]. Male and female meiosis differ on molecular level [58] and the genetic bases of recombination rate also varies between sexes [21]. Thus, recombination can only be understood by studying both male and female meiosis. However, experimental data on a molecular level is scarce beyond few model organisms and thus comparative genomics may help by linking both the similarities and differences of recombination to other genomic features at different scales of genome organization.

We chose to compare recombination between fish species due to their diversity and availability of suitable data. Fish have spread to all kinds of aquatic environments across the world and comprise approximately half of all extant vertebrate species [20, 31]. Fish allow studying both the variation of the total number of recombinations and their positioning within genomes. First, fish have highly varying levels of heterochiasmy. Consistent with the findings in other groups, majority of studies have reported female fish to have higher recombination rate when there is a difference between sexes but also the converse has been reported for many species [14]. Fish also share the trend observed accross many taxa that male recombination is often more concentrated towards the ends of the chromosomes than female recombination [39]. Second, most but not all teleosts appear to have lost the complete PRDM9 needed to direct crossovers [4, 12] and thus allow studying the positioning of crossovers both with and without PRDM9.

Variation of recombination across species has been studied by comparing their linkage maps [45]. In linkage mapping the inheritance of alleles from parents to their offspring is followed at a large number of genetic marker positions across the genome. Comparing the allele patterns at consecutive marker positions then allows inferring when a crossover has occurred between the markers. When done for a large number of offspring and markers, this provides a map of genetic distances between markers (measured as centimorgans, cM, 1% observed recombination frequency between the markers) across the genome.

Linkage mapping readily allows comparing sexes as separate maps can be constructed for male and female parents. Recently, methods based on sequencing either population samples or gametes have become popular alternatives to study recombination [34]. As measuring association of alleles from a population sample only allows inferring rates of recombinations that have accumulated over many generations, it does not provide sex-specific information. Direct sequencing of gametes is often limited by the availability of suitable samples and so far has been mainly used to study male recombination.

Despite the large number of linkage maps generated for individual species, quantitative comparison of species by comparing linkage maps generated in different studies has been challenging. First, many studies have not made the generated linkage maps available [18]. Thus, the maps have typically been compared on a high level such as how much recombination is concentrated towards chromosome ends [18, 39], prohibiting for example the comparison of local recombination rates with other genomic features. On the the other hand, if sex-averaged maps are compared [9], varying differences between the sexes in the compared species may confuse the results. Second, the true differences between species can be obscured by non-biological differences between the studies [14] and thus stringent filtering and corrections need to be applied before comparisons can be made [9, 14].

Recently, combining traditional linkage mapping with highly parallel sequencing has allowed the generation of sex-specific high-resolution recombination landscapes. We set out to generate the first comparative set of sex-specific linkage maps covering a wide variety of species from a large taxonomic group. To avoid problems caused by differences in data handling, we decided to start from raw sequencing data and generate all linkage maps using the same computational pipeline.

The main limitation of linkage mapping is that it requires genotyping a large pedigree, at least dozens but preferably hundreds of individuals from one or more nuclear families. Thus, for most vertebrate groups, large data sets are available only for a few domesticated or model species. Fortunately, many fish species generate large numbers of offspring. Thus, we took advantage of the availability of suitable data sets and were able to generate high-density linkage maps for 40 diverse fish species.

## Results

### New linkage maps allow comparisons across fishes

We searched and curated published fish data sets that could be suitable for linkage mapping from literature and data repositories. We concentrated on sequence-based studies for which the read-level data was publicly available (see Methods for details). To enable quantitative comparison between species, we processed the data from the different studies through the same computational pipeline. We were able to generate sex-specific linkage maps for 40 fish species (Fig. 1, Supplementary Table 1). The number of offspring ranged from 78 to 1,738 (median 145) and the number of markers from 1,837 to 2,268,031 (median 34,068) (Supplementary Table 2). Despite the large range, the length of the linkage map did not depend on the number of markers used (Pearson correlation coefficient between the number of markers and the sex-averaged map length -0.01).

**Fig. 1.**
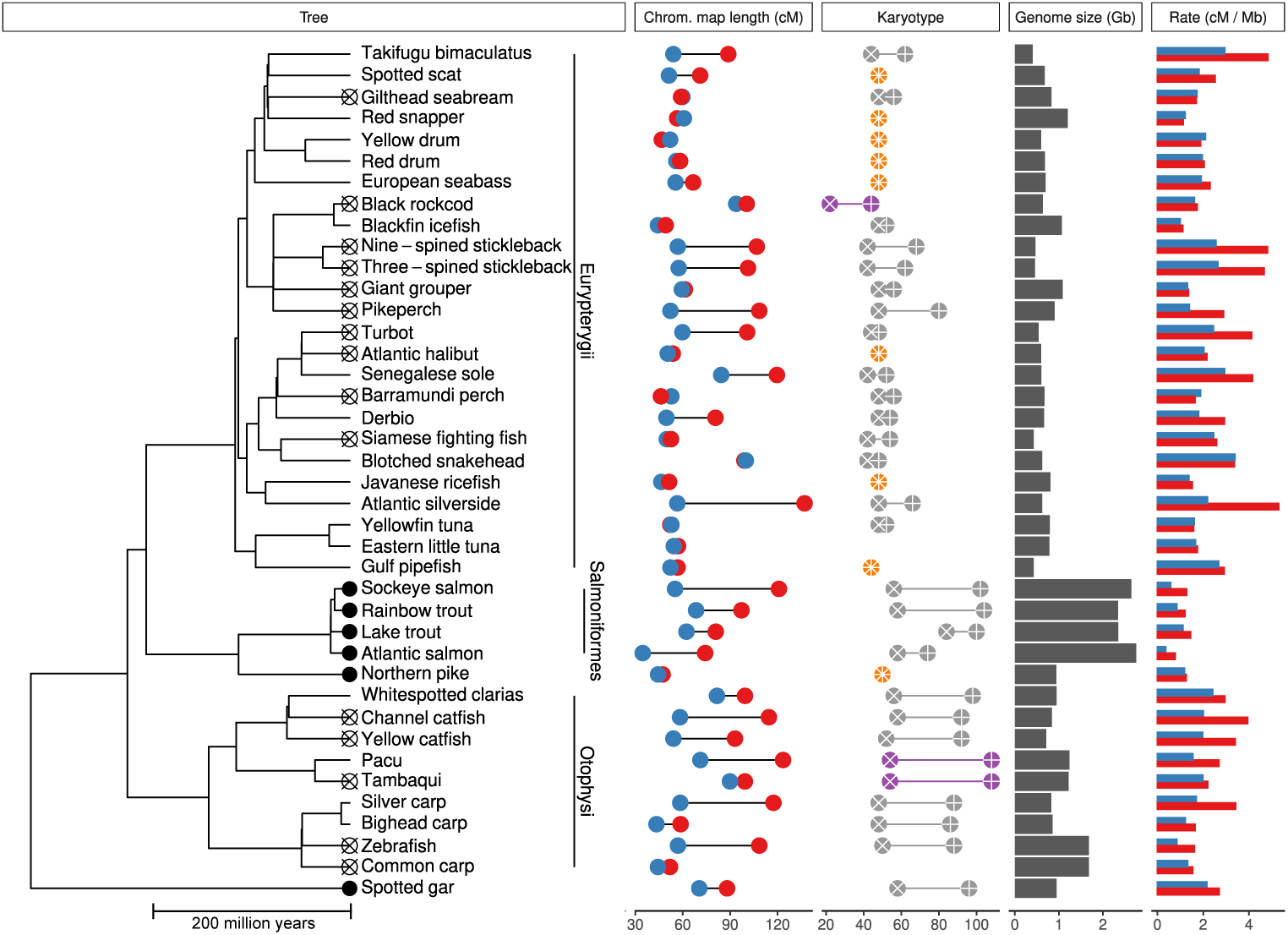
Fish species for which new linkage maps were created. The species that have retained or lost a complete PRDM9 gene according to [12] are labeled on the phylogenetic tree with filled and crossed circles, respectively. The major taxonomic groups covered by more than one species are shown next to the tree (see text for details). Average male (blue circles) and female (red circles) linkage map lengths per chromosome as well as diploid chromosome (cross) and chromosome arm (plus) numbers are shown for each species (karyotype not known for eastern little tuna and not shown for tetraploid common carp). The fully acrocentric (orange) and metacentric (purple) species are highlighted. Genome sizes as well as male (blue bars) and female (red bars) recombination rates are shown.

To relate recombination to the genome structure, we also collected available karyotype information on the mapped fish species. Unfortunately, the exact chromosomal positions of the centromeres are not yet available for most species as centromeric sequences are difficult to assemble. However, for 38 out of the 39 diploid species the number of acrocentric (centromere close to one end of the chromosome) and metacentric (centromere close to the middle of the chromosome) was available or could be inferred from close relatives (see Methods for details).

The species in the linkage map collection belong to the main extant fish lineage, teleosts [20, 31], except the outgroup species, spotted gar, that has been termed a living fossil [11] (Figure 1). Teleost fishes have two small and two large subdivisions Euteleostei and Otocephala. A majority of the fish (25 species in 10 different orders) in the collection belong to Euteleostei subgroup Eurypterygii that includes 60% of all teleost species many of which have karyotypes dominated by acrocentric chromosomes [55]. The rest of the Eutelostei species in the collection belong to orders Salmoniformes (4 species) and Esociformes (northern pike) that have retained a complete PRDM9 gene [12]. The Salmoniformes have long genomes and varied karyotypes compared to most Euteleostei because their ancestor had a whole-genome duplication event after the branching from the common ancestor with Esociformes [38]. The rest of the species (9 species, 3 orders) in the collection belong to the Series Otophysi, a subgroup of Otocephala, that includes 32% of teleost species, characterized by karyotypes with high arm numbers [55]. In summary, the collection covers a diverse set species across the teleosts (15 different orders).

The species averages of sex-averaged chromosome map lengths varied from 46.9 cM (blackfin icefish) to 102 cM (senegalese sole). For some species, the average male chromosome map length was considerably under 50 cM but this is probably due to difficulties to reliably detect crossovers very close to the chromosome ends based on limited data. Thus, as half of all crossover events are observed in the offspring, a typical fish chromosome pair has on average one to two crossovers in a meiosis.

### Higher recombination rate in females is associated with mixed karyotype

In general, the sexes either had similar recombination rates or the females had a higher rate. The differences between species were mainly driven by female recombination (female average chromosome map length 83.0 cM, coefficient of variation 0.39 vs. male 59.1 cM and 0.32, respectively).

The species with substantially higher female rate were scattered across the phylogenetic tree without any obvious pattern (Fig. 1). However, combining recombination rates and karyotypes revealed a strong general pattern (Fig. 2). In both extremes of the karyotype spectrum, i.e. all chromosomes either acro- or metacentric, both sexes had approximately one recombination event per chromosome arm, i.e. one per chromosome if all were acrocentric and two if all were metacentric, respectively (Fig. 2a,b,c). In contrast, the difference between the sexes was largest when the minority karyotype accounted for more one third of all chromosomes. Typically in these species females had two recombination events per chromosome on average whereas males had close to one, respectively. In other words, if the species carried both types of chromosomes, females were typically closer to the metacentric (two events per chromosome) and males to the acrocentric (one event per chromosome) pattern, respectively. Thus, recombination was not proportional to the number of chromosome arms in either of the sexes (except near the ends of the spectrum).

**Fig. 2.**
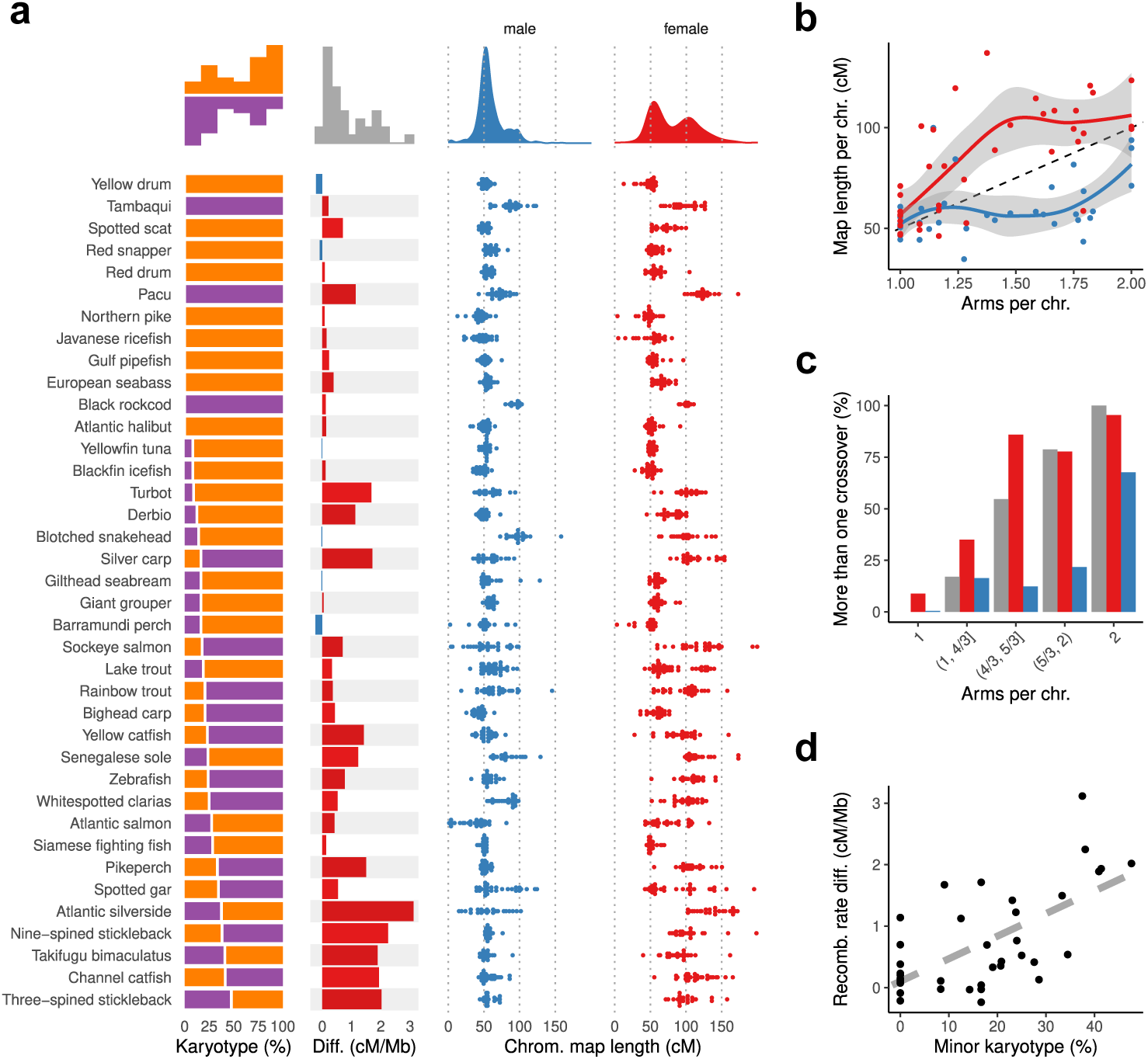
Fish karyotypes and global recombination rates. **a**, Left panel, the percent of acrocentric (orange) and metacentric (purple) chromosomes of all chromosomes. Middle, the difference between female and male recombination rates, the species with higher male and female rates are colored blue and red, respectively. Right, chromosome linkage map lengths for males (left, blue) and females (right, red). The species are ordered according to the percentage of chromosomes that are of the minority type (acrocentric or metacentric). Average histogram (left, middle) or smoothed density (right) calculated over all species is shown over each panel. **b**, The sex-specific (female red, male blue) average chromosome map length (y-axis) is plotted against the average number of arms per chromosome (x-axis) for each species. Loess fits (female red, male blue line) and their 95% confidence intervals (shaded areas) are shown. The black dashed line represents one recombination event per chromosome arm. **c**, Number of chromosomes with more than one crossover compared to expected. A chromosome was classified to likely have more than one crossover if the map length was over 75 cM. The percentage of such chromosomes (y-axis) in females (red) and males (blue) is compared to expected (gray) for five groups of species defined based on the arms to chromosome ratio (x-axis). The expected number for each group is calculated assuming one crossover per chromosome arm. **d**, Karyotype predicts recombination rate difference between sexes. For each species (n=38), the difference between female and male recombination rates is plotted as a function of the minor karyotype percentage. The gray line shows the fit of the PGLS model.

The difference between male and female recombination rates correlated strongly with the karyotype also when the evolutionary history of the fish species was taken into account (Fig. 2d, p-value *<* 10*^−^*^5^, phylogenetic generalised least squares model, see Methods for details). The phylogenetic signal was estimated to be weak (maximum likelihood estimate *λ* = 0, 95% confidence interval upper limit 0.53), indicating a weak relation between the trait values and the phylogeny. This was expected as the species with the largest differences between sexes but similarly mixed karyotypes such as atlantic silverside, sticklebacks, and channel catfish, represented distant branches of the phylogenetic tree (Fig. 1).

### Recombination and sequence features

Next we wanted to see which sequence features associated with recombination. First, we calculated the Pearson correlation coefficients (from now on correlations) between recombination rates and all ten dinucleotides over 32 windows per chromosome excluding repeats (Fig. 3). Correlation patterns varied both between species and sexes of the same species. Broadly, the dinucleotides lacking C and G were negatively associated with recombination but even this trend was reversed in some cases, most prominently in spotted scat females. The correlation patterns of the GC-containing dinucleodides were more complicated. In males, majority of species exhibited the same correlation pattern to a varying degree (Fig. 3a). In this pattern, the CpG correlation is coupled with the CpA:TpG anticorrelation. This pattern can emerge if the regions of high recombination rate are hypomethylated in the germ line cells compared to the rest of the genome as methylated CpG:s are prone to mutate to TpG:s [49]. In contrast, all four Salmoniformes had a distinct male correlation pattern that had no apparent relation to GC or CpG content and was different from all other species, including spotted gar and northern pike that also have a complete PRDM9 gene.

**Fig. 3.**
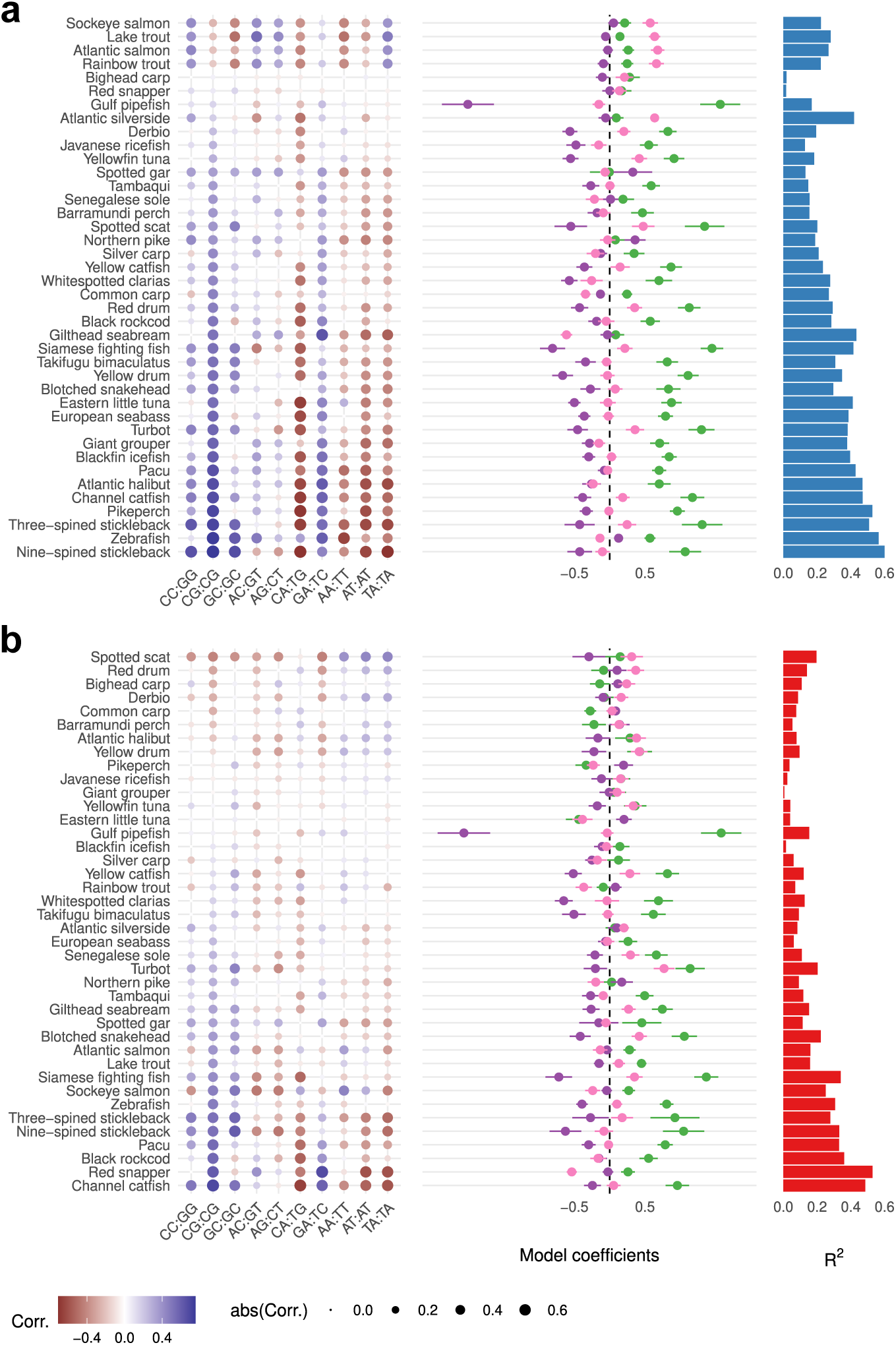
Association of recombination and dinucleotide content of the (repeat masked) genome. **a**, Left, correlations between dinucleotides and male recombination. The species (rows) are ordered according to the correlation of the most highly correlated dinucleotide CpG (CG:CG). Middle, the coefficients of the linear models predicting recombination rates using CpG (green) and CG (dark purple) contents as well as DNA bending (light purple). Right, the variance explained (*R*^2^) by the multivariate linear models. **b**, As in **a** but for female recombination.

Next, we generated multivariate linear models predicting recombination in each species and sex separately to see how well three sequence characteristics linked to biological processes, GC and CpG content as well as DNA bendability, could explain the observed dinucleotide patterns. We calculated predicted DNA bendability because it is generally independent of GC content and is associated with many central biological processes [5]. Also, some of the components of the recombination machinery favor bent DNA [58]. In most species, the CpG content was clearly the dominating factor of three in both sexes, especially in those species in which the models had the highest predictive power. In contrast, GC content was typically not positively associated with recombination when the two other factors were taken into account. In few cases such as male Salmoniformes and atlantic silverside, DNA bending had the largest effect. In general, the tested sequence features had less predictive power in females than in males (Fig. 3b). Overall, CpG content was clearly the most prominent feature linked to biological processes but there were also features such as the high correlation of GpA:TpC in males that could not be fully explained.

Our collection contained whole-genome data for four species. For these species, we were able to look at the sequence features in the immediate vicinity of the crossovers (Extended Data fig. 1). We compared the abundance of each dinucleotide to its whole-genome average at each position relative to the crossover and contrasted this to the average computed over 20 kb windows around the crossovers. Mostly the regions around crossovers showed similar trends as the whole-genome correlation analysis (Fig. 3), but there were also notable exceptions. For example, in pikeperch males the immediate neighborhood of crossovers had a different pattern than the broader flanking region. Generally, CpG was again the mostly highly enriched dinucleotide. In males of three species (derbio, pikeperch, and takifugu bimaculatus) the CpG content close to the crossovers was over forty percent higher than the whole-genome average. In summary, recombination associates with sequence features both in broad (from tens of kilobases to megabases) and fine (kilobase) scales.

Transposable elements and other repeat elements have the ability to rapidly reshape the genomic landscape of a species. Thus, we annotated repeats for all our genomes and compared the repeat content to the recombination rate in the 32 windows per chromosome. Our repeat annotation is incomplete, as the genome assemblies lack some of the highly repetitive regions such as centromeres that are difficult to assemble. Also, a large fraction of the *de novo* repeats found from the genomes could be classified (class ”Unknown”). Nevertheless, the comparison revealed large differences between the species and even between sexes of the same species (Extended data figures 2 and 3). The correlations between recombination rate and repeat coverage ranged from strongly negative (*−*0.51 in Siamese fighting fish males) to strongly positive (0.52 in pikeperch males). The associations were not driven by the total fraction of the genome annotated as repeats (male and female correlations 0.16 and 0.19, respectively, p-values *>* 0.2). Most transposable element classes could have both positive and negative association and only few classes like TcMar-Tc1 in males showed a clear negative association across majority of species. On the other hand, simple repeats and low complexity regions were positively associated with recombination in males across species.

### Chromosome recombination rate and sequence features

Next we investigated whether simple sequence features alone could partly explain chromosome level recombination rates (Fig. 4). Chromosome lengths varied by an order of magnitude, yet they typically had on average one to two and one to three crossovers per chromosome in males and females, respectively, regardless of their size (Fig. 4a). Thus, short chromosomes had higher recombination rates (cM/Mb) than the long ones.

**Fig. 4.**
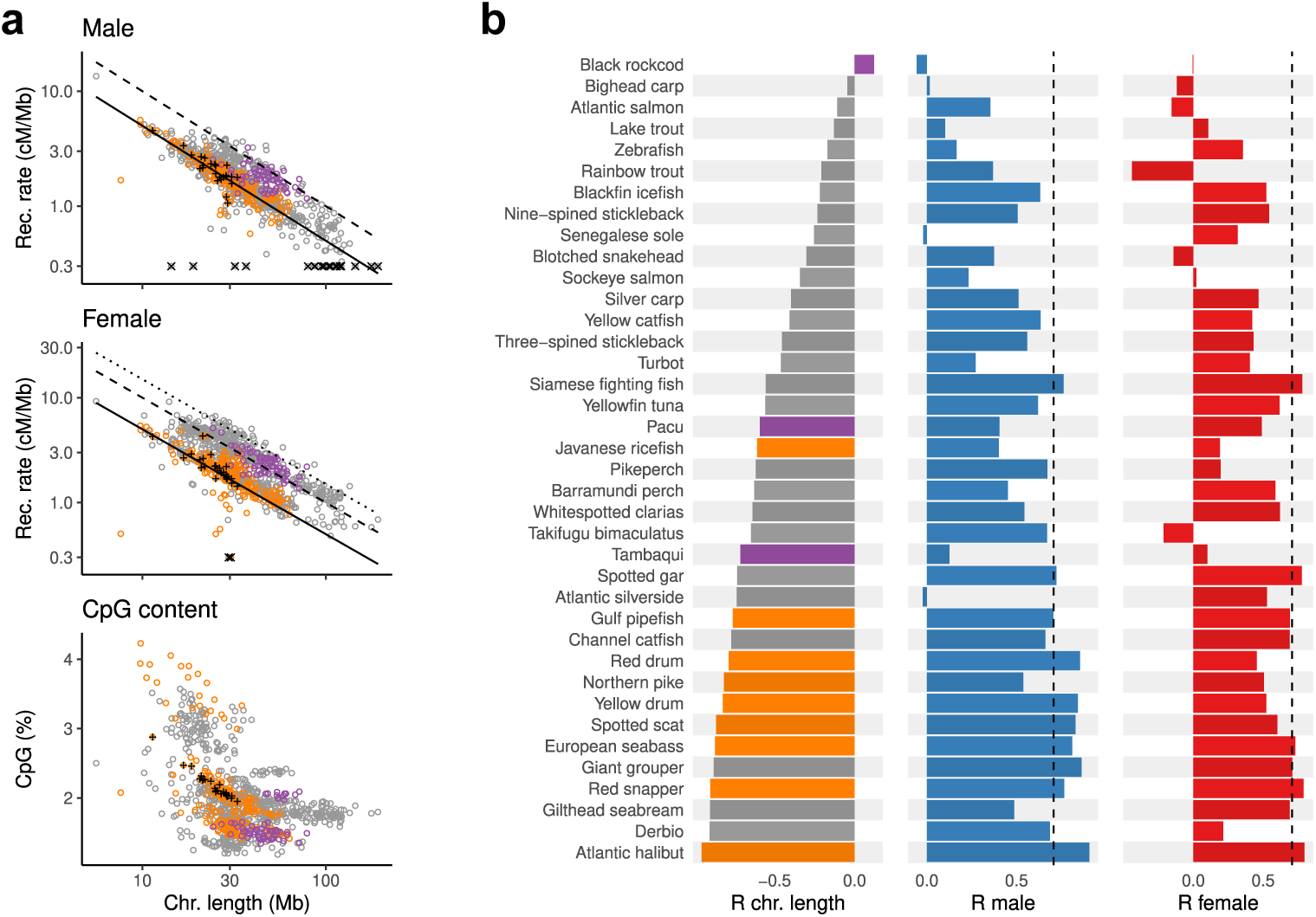
Variation of recombination rate between chromosomes. **a**, The male (top), female (middle) recombination rates (in logarithmic scales) and CpG% (bottom) are shown for each chromosome as a function of its size (x-axis, logarithmic scale). The solid, dashed, and dotted lines represent recombination rates equal to one, two and three recombination events per chromosome, respectively. As an example, atlantic halibut chromosomes are marked by a plus in all panels. The rates smaller than 0.3 cM/Mb are shown at 0.3 and marked with a cross. The karyotype of the species is shown by color, mixed (bottom layer, gray), all acrocentric (middle, orange), and all metacentric (top, purple). **b**, The correlation of CpG content and chromosome length (left; coloring according to karyotypes as in a), male (middle), and female recombination (right panel) for each species. All correlations are between relative values obtained by dividing the chromosome value (length, CpG content, recombination rate) by its average over all chromosomes of the species. The dashed black line marks correlation that explains half of the variation between chromosomes.

Generally, the short fish chromosomes had a higher CpG content than the long ones (Fig. 4a). Thus, also within a genome, a short (or long) length of a chromosome compared to the others could partly be compensated by the high (low) CpG content. If this was the case, we would expect the relative CpG contents of the chromosomes to partly explain their relative recombination rates (compared to the within species averages). Indeed, in males in eleven and in females in seven species, respectively, the CpG content explained more than half of the recombination rate variation between the chromosomes (Fig. 4b). The effect was particularly strong in fully acrocentric species. In contrast, there was no effect in some of the species that have partly or fully metacentric karyotype. In summary, processes linked to CpG content not only shape the local recombination landscape but can also contribute to short and long chromosomes having approximately the same number of crossovers.

### Distribution of crossovers along chromosomes

In many species, the positioning of crossovers within chromosomes also varied drastically between the sexes. We measured the similarity of crossover distributions between sexes by correlation of sex-specific recombination rates in the same 32 windows per chromosome as used above. The correlations ranged from highly positive to negative (Fig. 5a,b, Extented Data fig. 4). A similar genome-wide recombination rate in males and females did not imply similar positioning of crossovers. In fact, the correlation between the local male and female rates was higher in species that had a large difference in genome-wide rate, but the association was not significant (Fig. 5c).

**Fig. 5.**
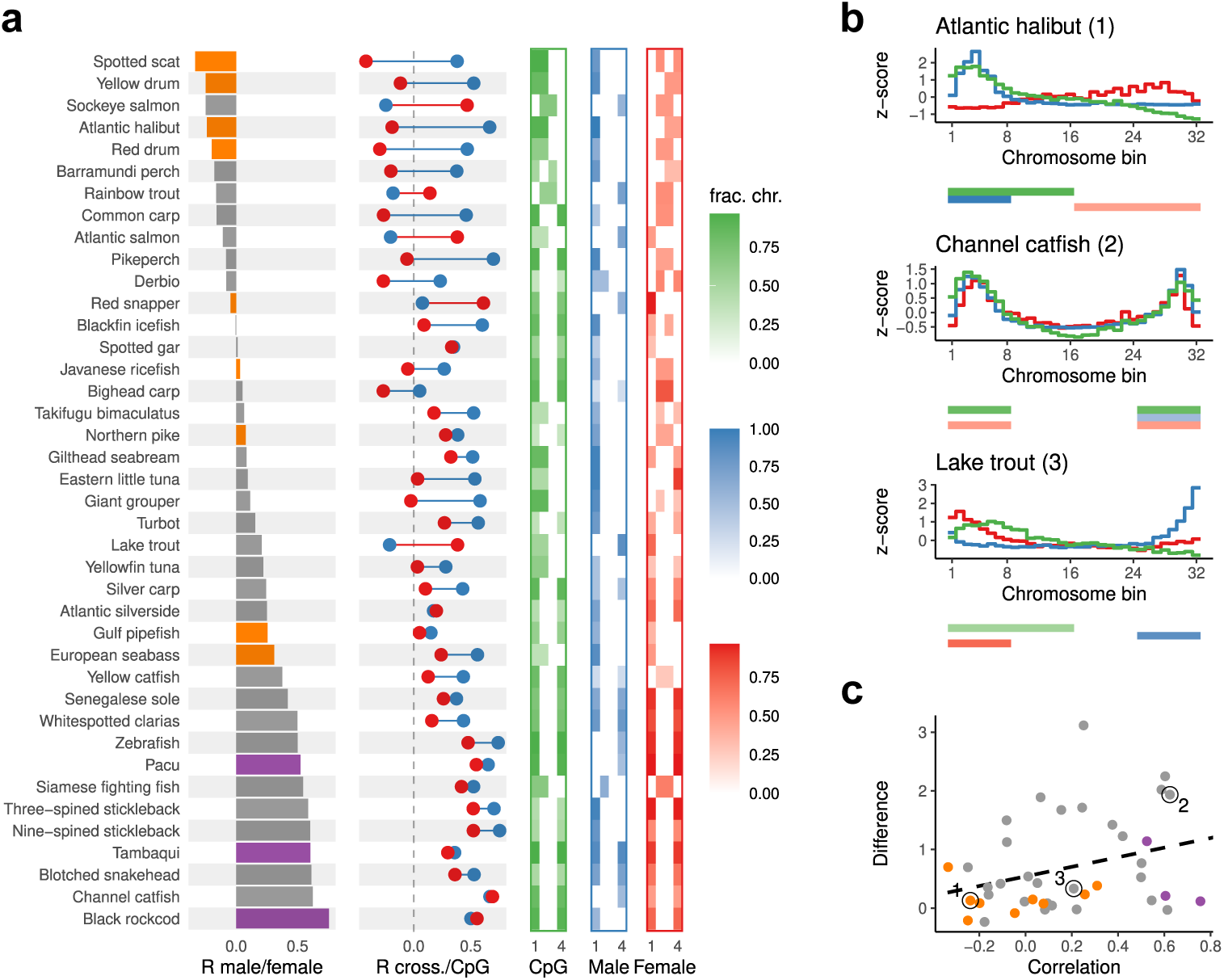
Local recombination rate and CpG content. The features where calculated in 32 windows per chromosome (repeat masked genome). **a**, The correlation between male and female recombination rates (left; coloring according to karyotypes as in Fig. 1), between recombination rates and CpG content (middle, female red, male blue) and the most common summary pattern for each feature (the darker the color the larger fraction of the chromosomes have the most common pattern shown; see the text for details). The species are ordered according to the correlation between the male and female recombination. **b**, Average male (blue) and female (red) recombination rates of the windows over all chromosomes plotted together with CpG content (green) in three example species (values have been normalized to z-scores). The corresponding summary pattern is shown below each plot. **c**, The genome-wide difference between the male and female recombination rates (y-axis, cM/Mb) plotted against the correlation of the local rates (x-axis), correlation 0.27, p-value 0.09. The example species are labeled with the numbers shown in b.

Whether the males and female crossovers occurred in the same or different regions of the genome was clearly related to their association with CpG content. In all the species in which male and female crossover distributions were highly similar, the recombination was also highly associated with CpG content in both sexes (Fig. 5a, bottom, and Fig. 5b, channel catfish). Conversely, if the distributions were highly different, the male recombination had high positive and female negative correlation with CpG content, respectively (Fig. 5a, top, and Fig. 5b, atlantic halibut), except for Salmoniformes that had the opposite pattern (see for example Fig. 5b, lake trout). In summary, the male and female recombinations were located in the same regions only if both were preferentially located in CpG-rich regions.

Next we wanted to visualize the broad-scale relationship between the positioning of recombination and CpG content along the chromosome. We first oriented each chromosome so that the half that had a higher CpG content was set first. Then we summarized the distribution of the feature (recombination or CpG) along each chromosome by dividing the chromosome to four equal-sized parts and marking as enriched the (one or two) quarters that accounted for more than half of the total amount of the feature in the chromosome. For each species we then visualized the enrichment pattern that accounted for the largest number of chromosomes (Fig. 5a).

The male recombination was typically concentrated towards the ends of the chromosomes. In most species, this was the case also in females but enrichment in the center was also observed. The sex differences in relation to the CpG content were clearly visible even on this coarse level. When male and female recombination anticorrelated (Fig. 5a, top), the CpG content alone oriented the chromosomes so that male recombination typically concentrated to the same (most species) or the opposite ends (Salmoniformes and red snapper). In contrast, in these species, the female recombination concentrated to the other parts of the chromosomes than the male recombination. Notably, this set included species such as yellow drum and atlantic halibut (Fig. 5b) that had no heterochiasmy (Fig. 2). On the other hand, if the male and female recombination were highly correlated, typically there was no clear orientation but both CpG content and recombination concentrated towards the ends of the chromosomes (Fig. 5a, bottom). In summary, CpG content was indicative of an orientation of whole chromosomes that affected also the high-level distribution of crossovers.

## Discussion

Studying the variation of recombination has been hampered by the lack of data collections that would allow quantitative comparisons across a wide range of species. Typically variation of recombination has been studied by directly comparing summary statistics of linkage maps generated in separate studies. However, linkage map construction is complicated by errors in the data as well as missing data. Differences in dealing with these uncertainties, both within the methods used to construct the linkage map [36, 54], as well as in the choices made during the map construction such as data filtering, can have a large effect on the result. As a result, non-biological differences between the studies can obscure or distort true biological differences between the species. For example, in a recent meta-study in which sex-specific linkage maps of 61 fish species across 13 orders were compared, a strong correlation was found between the number of markers used in the study and resulting linkage map length [14]. We believed that maps that had been constructed in a consistent manner throughout would be more comparable and thus chose to construct all the compared maps using the same computational pipeline. Accordingly, in our collection the number of markers does not correlate with the linkage map length (correlation -0.01) even though the numbers markers used span from few thousand to millions.

Our collection covers 40 fish species across 15 different orders. As in many other groups, fish chromosomes typically had one to three crossovers per meiosis irrespective of their size. In mammals, it has been observed that sex-averaged recombination is roughly proportional to the number of chromosome arms and therefore one crossover per arm was proposed as a general rule linking karyotype and recombination (see [51] and the references therein). Our sex-specific maps show that at least in fishes neither male nor female recombination follows this rule when there are both acrocentric and metacentric chromosomes (Fig. 2). However, since males and females deviate from the rule to the opposite directions, i.e. females have more and males less crossovers than expected by the rule, their average can be close to one crossover per arm.

To our knowledge, no trait has quantitatively been shown to associate with the level of heterochiasmy. Here we showed that a surprisingly simple measure characterizing the karyotype of the species can partly explain the variability of heterochiasmy among fishes (Fig. 2d). This is surprising given that the classification of the chromosomes to two distinct types is clearly a simplification. The association between karyotype and heterochiasmy was not due to evolutionary relatedness of the species. Thus, it seems that heterochiasmy has not evolved by small gradual changes but is more likely labile on the time scale of teleost evolution. These facts would be consistent with a scenario in which recombination needs to adapt over and over again to changes in karyotype. Notably, massive changes in karyotype can occur rapidly [10].

Heterochiasmy was mainly driven by the increase of the female recombination in the species that had mixed karyotype compared to the fully acrocentric species while the male recombination was more stable. Thus, it seems that the forces driving the changes in global recombination rate mainly act during the female meiosis. One force exclusive to only one sex is female meiotic drive, biased segregation of homologous chromosomes either to polar body or egg, probably driven by differences in centromere strength [48]. As chromosome fusions and fissions alter centromere strengh, female meiotic drive is thought to have a profound effect on karyotype evolution [7, 13] and explain why both in mammals [50] and fish [29] karyotypes are biased towards all or almost all chromosomes having the same morphology i.e either acrocentric or metacentric. Evolutionary theory predicts that female meiotic drive leads to rapidly evolving female recombination rates [8, 39] but directly detecting such effects remains elusive. Our findings support meiotic drive as a potential cause of heterochiasmy.

As the differences in genome-wide and local recombination rates between sexes were unrelated (Fig. 5c), karyotype changes do not seem to be associated with recombination rate changes in particular genomic regions but rather with an adjustment of the total rate affecting the whole genome. Mammalian studies have revealed genetic loci that have large and often sex-specific effects on recombination rate [21]. Thus, rapid change of the genome-wide recombination rate is possible. One mechanisms allowing such global adjustment is the modulation of chromosomal axis length [58].

Lately, much of the attention has concentrated on recombination hotspots, narrow regions of high recombination rate. Consistent with other studies on species that lack complete PRDM9, we observed enrichment of CpG:s close to crossovers that may indicate that recombination locally favors unmethylated DNA and open chromatin in these species (Extented Data fig. 1). In takifugu bimaculatus, all only GC-containing dinucleotides were enriched and all only AT-containing dinucleotides were depleted close to crossovers, which is consistent with GC-biased gene conversion occurring in the same narrow regions as crossovers.

It has been suggested that GC-biased gene conversion could be a major driver of genome base composition in many vertebrate groups including mammals and reptiles [15, 30]. Within the species studied here, GC content significantly associated with recombination in genome-scale only in few species when other sequence features were taken into account (Fig. 3). Thus, we did not find evidence for GC-biased gene conversion being a common driver of genome-scale genome composition in fishes. However, comparing mutation spectra of different genomic regions is needed to further understand the links between recombination and other drivers of genome base composition [6].

Instead of general GC content, CpG content was the dominating sequence feature explaining the broad-scale variation of the recombination rate both between and within chromosomes. Particularly in males, CpG enrichment revealed the orientations of the whole chromosomes so that recombination typically concentrated to the same ends as CpG content (Fig. 5). Furthermore, CpG content alone explained a large fraction recombination rate variation between chromosomes (Fig 4b). Which mechanisms could cause such broad-scale associations that are often sex-specific? Two recent findings in mammals point to the potential role of replication timing. First, in male germline cells recombination is linked to replication timing [35]. Replication speeds of chromosomes are predictive of their crossover rates. In addition, subtelomeric regions replicate early and have an elevated GC content as well as recombination rate compared to the other parts of the chromosomes. Second, the high mutation rate of methylated CpG:s is not only caused by spontaneous deamination as previously thought but replication errors are a major contributor as well [49]. Morever, the CpG mutations are more effectively repaired in active chromatin and early replicating DNA, and the effect is not fully explained by the methylation levels of the regions [41]. Therefore, the replication errors may lead to fewer CpG mutations in the early than in the late replicating regions. Taken together, these observations are consistent with the male meiotic recombination occurring in early replicating regions that are generally less methylated, more active, and more efficiently repaired than late replicating regions leading to the broad association between recombination rate and CpG content we observed in fish. An intriguing question is why in some fish, the recombination in females follows CpG content like in males but in others, this link is totally absent or even reversed.

Based on few example species per each large taxonomic group, it has been assessed that recombination is typically negatively correlated with transposable elements [23]. Importantly, the centromers and their flanking pericentromeric regions, in which recombination is typically suppressed, consist of highly repetitive sequences. These regions are clearly underrepresented in our analysis because most fish genome assemblies currently lack the most difficult regions. However, we showed that generally the association between recombination and repeats varies drastically between species (Extented Data fig. 2 and 3) This may reflect the complicated dynamics between transposable element insertion and inactivation and chromatin landscape [47].

In summary, we have presented the first collection of consistently generated sex-specific recombination landscapes that highlights both similarities and differences between species. Our results emphasize the importance of obtaining the landscapes for both sexes, when possible, because of the differences in male and female meiosis. We expect the collection to promote further studies to elucidate how recombination interacts with other fundamental biological processes and evolutionary forces on different time scales to shape the genomes of sexually reproducing species.

## Methods

### Data collection

Literature and data repositories were searched for fish data sets suitable for high-resolution linkage mapping. Only sequencing based studies (RAD-seq, WGS or similar) that had pedigree and raw sequencing data as well as a reference genome of the species publicly available were considered. Studies based on hybrid offspring of two species were excluded. In addition, the pedigree had to be such that it enabled generating both male and female linkage maps. No hard limits were set for the sample size but generally studies that had in total at least close to 90 offspring in one or several (full- or half-sib) families were considered. For nine-spined stickleback, a map constructed of samples from multiple studies was used (Rastas, Kivioja, manuscript in preparation). The species were identified by their scientific names in NCBI taxonomy (Supplementary Table 1) and their phylogeny and divergence times were retrieved from TimeTree web server [24] (date of the download May 7, 2024) and visualized using R package ggtree (version 3.12) [56].

Raw sequence data and reference genomes were downloaded from the public data repositories (NCBI, ENA, CNGBdb), see Supplementary Table 1. Sample meta-data was collected from the databases and the original publications. If during data processing discrepancies were detected between the original publication, sample information and data, other independent information was searched to resolve the issue and if no such information was found, the study was not included in the collection (see the details below).

### Karyotype information

The majority of karyotype information was taken from the collection by Arai [2] (as summarized by Yoshida and Kitano [55] or directly from the book). For the species not covered in the collection, information was collected directly from primary publications (Supplementary Table 3). Following Yoshida and Kitano [55], we used the fundamental numbers (NF1) as the arm numbers. This numbering considers metacentric and submetacentric as two-armed and subtelocentric and acrocentric as one-armed chromosomes, respectively. For simplicity, we refer to the two types simply as meta-centric and acrocentric and calculated their numbers from the fundamental number of each species. (Other names have been used e.g. mono- and bi-armed [1] or bi- and monobrachial [7].)

For two species for which karyotype could not be found, red snapper and gulf pipefish, it was inferred based on the genus as in both cases all available species of the same genus had identical karyotypes. Tetraploid common carp was excluded from the karyotype comparison and no karyotype was found for a species in the same Genus as eastern little tuna.

### Linkage map construction

The linkage maps were generated using Lep-MAP3 [36] and Lep-Anchor [37] software. The workflows used are available at GitHub (https://github.com/kivioja/lep-pipe/). The raw sequencing reads were mapped to the reference genome using bwa mem or bwa-mem2. Duplicate reads were removed from Pikeperch high-coverage WGS data. The sequence coverage of the individuals were calculated using samtools stats and samples that had low coverage were discarded. Low coverage samples were identified manually for each species as samples that had a very low coverage relative to the rest of samples used for the same linkage map caused problems even if their absolute size was adequate. Variant calling was done using the mpileup command of the samtools program (version 1.13) and Lep-MAP3 module Pileup2Likelihoods as described in Lep-MAP3 documentation.

Pedigree information was validated using Lep-MAP3 IBD module. If the general family structure was unclear, the whole study was discarded. If family structure was clear but some individuals did not have the expected familial relationships, those samples were discarded but the study was kept. The sexes of the parents were taken from the supplied meta-data except for the following cases. The channel catfish parent sexes were swapped compared to the annotation based on the sex-linked marker given in the linkage data publication [57]. The resulting longer female map compared to male map was consistent with array-based linkage maps for the same species [26]. Turbot parent sexes were swapped compared to the annotation so that female map was longer than male, consistent with the original study [27] based on the same data and an independent study [28]. Takifugu bimaculatus parent sexes were swapped based on two sex-linked SNPs from another publication [53]. The resulting longer female map was consistent with the map given in the linkage data publication [40]. Derbio parent sexes were assigned based on the sex determining SNP given in the linkage data publication [17].

The linkage mapping was mostly automated. First, Lep-MAP3 modules Parent-Call2 and Filtering were used to call parent genotypes and to filter problematic variants. Additionally, if consecutive physical positions in the reference genome had variants, only the first one was kept. After running Lep-MAP3 module SeparateChromosomes2 with a range of logarithm of the odds (LOD) score limits, the results were manually checked and such LOD score limits were chosen that the number of major linkage groups matched the chromosome number of the species. If no such LOD score limits were found, the study was discarded. The linkage groups containing the largest numbers of markers were retained so that they number matched the number of chromosomes. Additional markers were added to the remaining linkage groups using the module JoinSingles2All. The markers were ordered by running the module OrderMarkers2 three times for each linkage group and taking the order that had the highest likelihood.

Next, a modified reference genome that was consistent with the linkage map was generated for each species using the Lep-Anchor tool. First, the contigs of the input reference genome were anchored to the linkage map by running Lep-Anchor iteratively using the wrapper script lepanchor wrapper2. Next, Lep-Anchor was allowed to detect and correct inconsistencies with the linkage map within contigs by running module PlaceAndOrientContigs with parameter findContigErrors=1. The resulting agp file for each species defined the new reference genome used in the rest of the data analysis. The coordinates of the markers in the linkage map were lifted over to the new reference genome and the final linkage map was generated by using the Lep-MAP3 module OrderMarkers2 to evaluate the linkage map in the order defined by the new genomic coordinates. Finally, the Lep-Anchor module EstimateRecombination was run for each linkage group to estimate the recombination density in a regular grid of 1024 points and to generate the corresponding Marey map.

### Phylogenetic generalised least squares

Phylogenetic generalised least squares (PGLS) model was fitted using the function pgls in R package caper version 1.0.3 [32]. The number of species used was 38 as for eastern little tuna the karyotype was not known and tetraploid common carp was excluded from the analysis. The fraction of chromosomes having the less common karyotype (acrocentric, metacentric) within the species was used as the predictor variable and the difference between female and male recombination rates as the response variable. The value of the parameter *λ* was optimized within default bounds using the maximum likelihood method, other parameters were fixed to their default values.

### Local recombination rates and sequence features

Local recombination rates were calculated for 32 windows per chromosome as follows. The Marey map that consisted of map lengths at 1024 points was divided into 32 windows (each consisting of 32 points) and the genetic length of each such window was calculated by subtracting the start point value of the window from its end point value. The local recombination rate was then calculated by dividing the genetic length of the window by its sequence length corrected by a scaling factor. Scaling was needed because in some of the species, a large fraction of contigs could not be assigned to the linkage map. Thus, the window sequence lengths were scaled by a factor calculated by dividing the total input genome assembly length by the total sequence length covered by the linkage map. (Consistently, the genome-wide recombination rates were calculated by dividing the total genetic map length by the total length of the input genome assembly.)

The sequence features were calculated in the same windows as the local recombination rates using the repeat masked genomes (see below). DNA sequence bendability was estimated using a model that takes into account dinucleotides and their pairs at variable distances [5]. The model parameters were set to the values trained with the unmethylated random library (Supplementary Table 10 of the article [5]). The bendability scores were calculated in 50 bp non-overlapping windows and the average of those scores was used for each recombination rate window.

Repeat models were trained for the input genome assembly of each species using RepeatModeler (versions 2.0.4-6) [43] and Dfam repeat database (version 3.7-8) [46]. The final assemblies generated by Lep-Anchor were annotated with the trained repeat models using RepeatMasker (versions 4.1.6-7) [44]. All the steps were done using Dfam TEtools Docker container.

## Acknowledgments.

The study was supported by Research Council of Finland (grant no. 343656 to PR). We wish to acknowledge CSC – IT Center for Science, Finland, for computational resources.

---

**Extended Data fig. 1.**
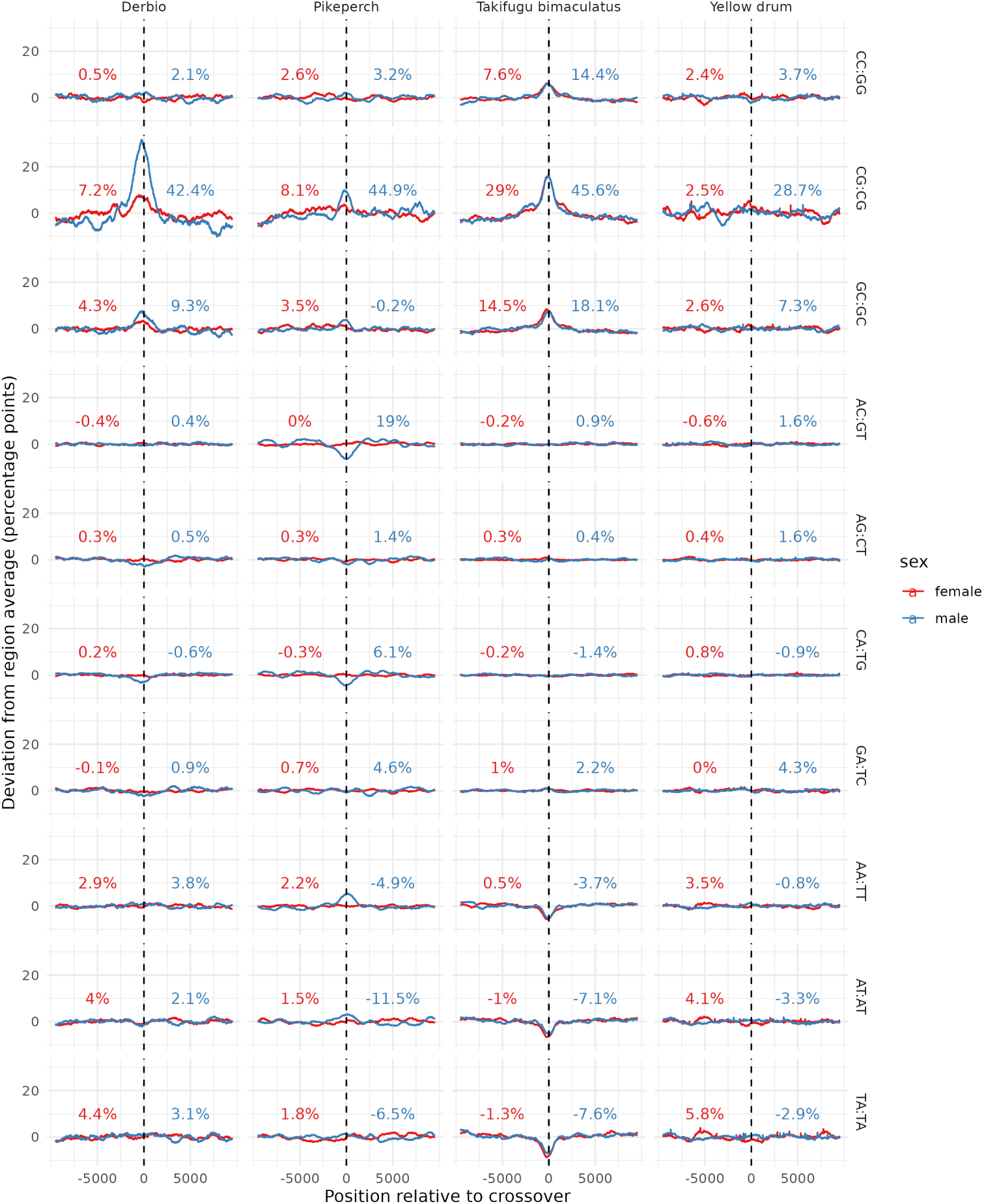
Dinucleotide content around crossovers in the four species that had whole-genome data. Each crossover region was centered at the middle point between the markers flanking the crossover (crossover was included only if the maximum distance between the flanking markers was less than 1 kb). Next the differences between the average abundances of dinucleotides (calculated using 1 kb sliding window) and their whole-genome averages were computed for each position around the middle point over all crossovers. The numbers (male blue, female red) show the maximum differences compared to the whole-genome averages. The lines (male blue, female red) show the deviations from the average differences computed over the whole 20 kb regions.

**Extended Data fig. 2.**
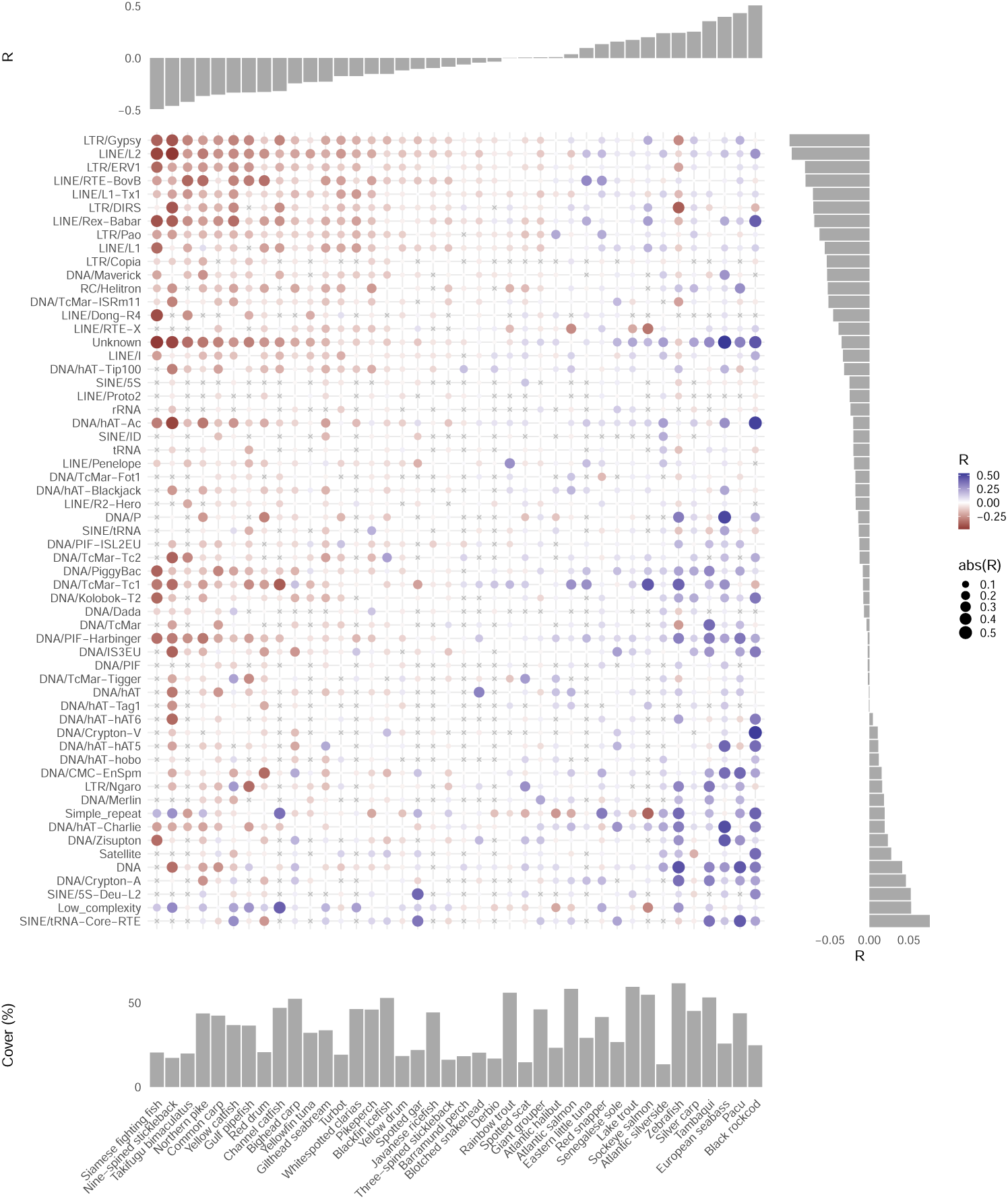
Association between female recombination and repeats. The heat map shows the Pearson correlations (R) between recombination rates and repeat class coverages calculated in 32 windows per chromosome (classes missing from a species are denoted by a cross; ”Unknown” contains all unclassified repeats). Only the repeat classes found in at least half of the species are shown. The species are ordered according the correlation between recombination rate and overall coverage by repeats in the same windows (shown in the top panel). The repeat classes are ordered according to the average correlation of each class over all species (shown on the right panel). The bottom panel shows the total fraction (%) covered by repeats calculated over all genomic windows.

**Extended Data fig. 3.**
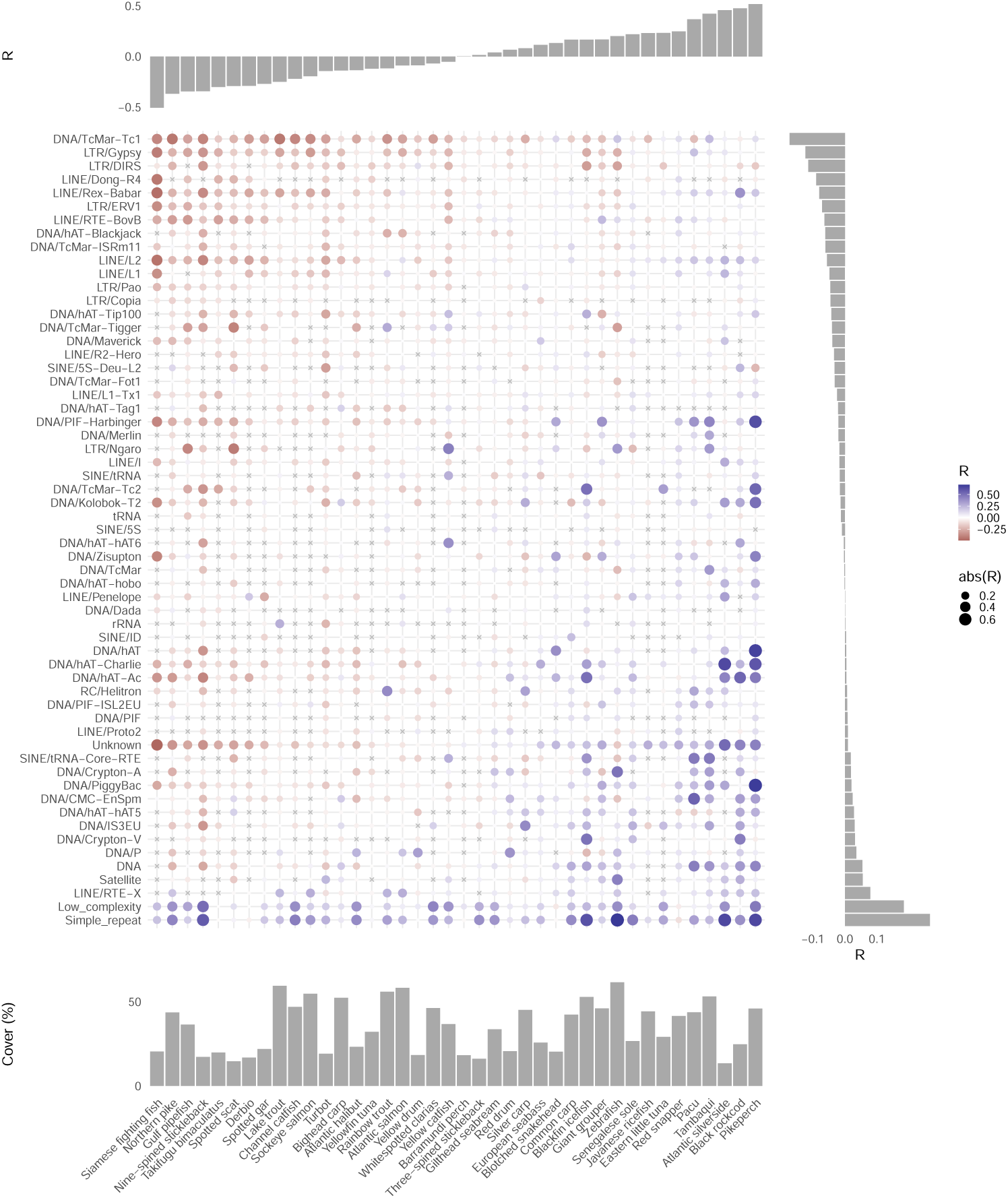
Association between male recombination and repeats. The heat map shows the Pearson correlations (R) between recombination rates and repeat class coverages calculated in 32 windows per chromosome (classes missing from a species are denoted by a cross; ”Unknown” contains all unclassified repeats). Only the repeat classes found in at least half of the species are shown. The species are ordered according the correlation between recombination rate and overall coverage by repeats in the same windows (shown in the top panel). The repeat classes are ordered according to the average correlation of each class over all species (shown on the right panel). The bottom panel shows the total fraction (%) covered by repeats calculated over all genomic windows.

**Extended Data fig. 4.**
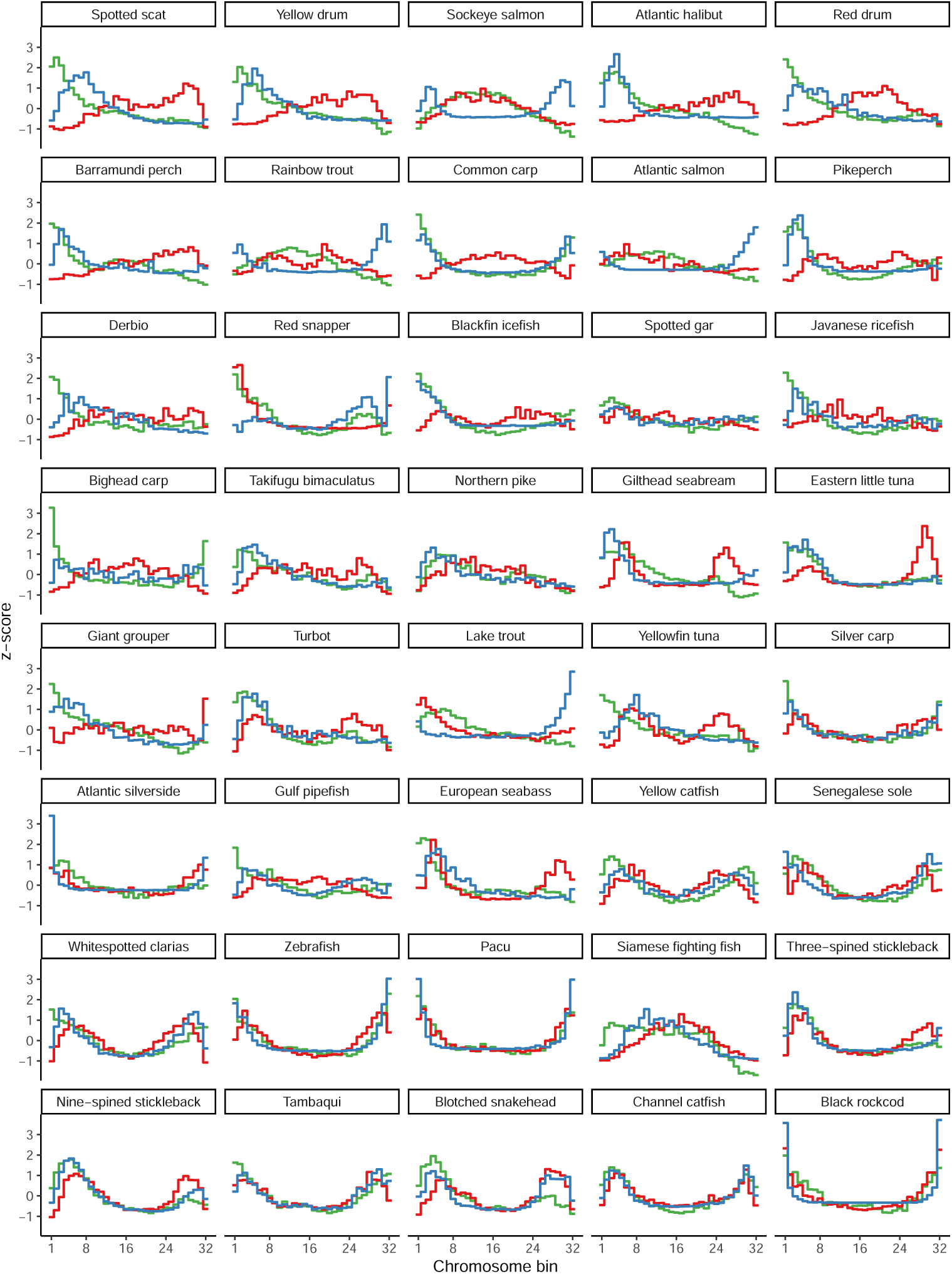
Average male (blue) and female (red) recombination rates of the 32 windows per chromosome averaged over all chromosomes plotted together with CpG content (green). All window values have been normalized to z-scores within species. The species are ordered according to the correlation between the male and female recombination.

